# Formate-driven isoprene reduction enables energetic independence in *Pelotomaculum schinkii*: Evidence from metatranscriptomic and functional analyses

**DOI:** 10.64898/2026.02.25.708027

**Authors:** Samikshya Giri, Anna E Rockwood, Ashley Logan, Sabrina Beckmann

**Author notes:** Conflict of interest declaration: The authors declare that they have NO affiliations with or involvement in any organization or entity with any financial interest in the subject matter or materials discussed in this manuscript. Author contributions: Samikshya Giri and Dr. Sabrina Beckmann contributed to designing and implementing the research, analyzing the results, and writing the manuscript. Dr. Sabrina Beckmann conceived the original idea and supervised the project. Anna E. Rockwood and Ashley Logan assisted with experimental work and data collection.

## Abstract

Isoprene, a volatile hydrocarbon emitted in vast quantities by terrestrial vegetation, can serve as a terminal electron acceptor in anoxic environments. Here, we investigated the molecular and physiological basis of isoprene reduction in an anaerobic enrichment culture highly enriched in Pelotomaculum. Substrate-exclusion assays revealed that formate is indispensable for isoprene reduction: cultures lacking formate showed no isoprene reduction, whereas formate alone sustained both reduction activity and the highest enrichment of Pelotomaculum. Metatranscriptomic analyses confirmed that *Pelotomaculum schinkii* was the most transcriptionally active species under isoprene-amended conditions, whereas *Sedimentibacter saalensis* was most enriched in unamended controls. Isoprene-treated cultures exhibited strong upregulation of [NiFe]-hydrogenase maturation factors (hypA/hybF, hypB), energy-conserving complexes (ATP synthase, H /Na - translocating pyrophosphatases), and transport systems (FeoB, ModBC), consistent with a formate-driven respiratory pathway. In contrast, genes for oxidative stress and metal detoxification were enriched in controls, suggesting physiological stress in the absence of isoprene.

Thermodynamic calculations further supported this mechanism, showing that formate oxidation coupled to isoprene reduction is highly favorable (ΔG°′ ≈ –135 kJ/mol), with an energy yield far exceeding that of formate-driven hydrogen production. Together, these findings support two mechanistic models: (i) direct isoprene reduction by group 4b [NiFe]-hydrogenase or (ii) indirect reduction via hydrogen transfer to an isrA-like oxidoreductase, analogous to *Acetobacterium wieringae* ISORED-2.

Our results demonstrate that *P. schinkii*, a metabolically versatile bacterium, couples formate oxidation to isoprene respiration, indicating that isoprene can serve as an alternative electron sink. This allows *P. schinkii to* escape obligate syntropy with methanogens and alters microbial activity and energy conservation in anoxic ecosystems.

## Introduction

Isoprene (C H) is the most abundantly emitted biogenic volatile organic compound (BVOC), with global annual fluxes estimated at nearly 500 Tg (Carrión et al., 2020; Sharkey et al., 2008). Terrestrial vegetation accounts for up to 90% of these emissions, making isoprene a dominant mediator of plant–atmosphere interactions (Carrión et al., 2020). Once in the atmosphere, isoprene readily reacts with hydroxyl radicals (OH), thereby altering oxidative capacity and extending the atmospheric residence time of greenhouse gases such as methane (CH) (Carrión et al., 2020; Collins et al., 2002). Consequently, isoprene plays an indirect but substantial role in regulating atmospheric chemistry and influencing global climate.

Beyond the atmosphere, soils serve as both a source and a sink for isoprene, harboring diverse microbial communities capable of producing and metabolizing this compound (Fall & Copley, 2000). Aerobic isoprene degradation has been well characterized, particularly in actinobacteria such as *Rhodococcus* strain AD45, which employs a multicomponent monooxygenase to initiate oxidative breakdown (Dawson et al., 2023; Sims et al., 2022; van Hylckama Vlieg et al., 1998, 2000). However, the anaerobic transformation of isoprene remains poorly understood. *Acetobacterium waeringae* is the only known anaerobe capable of reducing isoprene to methylbutenes, likely using it as a terminal electron acceptor to conserve energy (Jin et al., 2022; Kronen et al., 2019, 2023). This reduction is predicted to be catalyzed by a novel oxidoreductase in *Acetobacterium wierinage,* encoded within a dedicated five-gene operon (isr cluster) that is distinct from previously characterized ene reductases (Kronen et al., 2023).

A homology-based search for *isrA*, the key gene in the isr operon of Acetobacterium wieringae strain ISORED, identified its closest homologs (77% amino acid similarity) in *Pelotomaculum* species, including *Pelotomaculum schinkii* (Kronen et al., 2023). Members of this genus are strictly anaerobic, Gram-positive, spore-forming bacteria in the family *Peptococcaceae*, phylum Firmicutes, best known for oxidizing propionate in syntrophic association with methanogens (de Bok et al., 2005; Imachi et al., 2002). *P. schinkii*, in particular, has long been regarded as a strictly syntrophic propionate oxidizer that depends on interspecies electron transfer with methanogens to sustain growth (de Bok et al., 2005; Hidalgo Ahumada et al., 2018). Obligate syntrophs such as *P. schinkii* typically display distinctive ecological traits: they co-occur with specific partners, rarely dominate mixed enrichment cultures from the environment unless their niche is strongly selected, and collapse in abundance when their electron-accepting partner is absent, highlighting their reliance on syntrophic interactions (Morris et al., 2013b).

The presence of isrA-like homologs in *P. schinkii* suggests that it may use isoprene as an alternative electron acceptor under anoxic conditions, potentially decoupling its metabolism from strict syntrophic dependence. In our earlier work, we enriched a soil sediment culture highly enriched in *P. schinkii* that actively reduced isoprene in the absence of methanogenic or sulfate-reducing partners. Building on these observations, the present study investigates the molecular basis of isoprene reduction and establishes a mechanistic link between formate oxidation and isoprene respiration.

Our findings broaden understanding of how obligate syntrophs can use alternative electron-accepting processes, revealing unexpected metabolic flexibility that reshapes their ecological roles and contributions to carbon cycling.

## Materials and Methods

### Chemicals and standards

Isoprene (99% purity, Sigma-Aldrich), 3-methyl-1-butene (≥95.0%, TCI), 2-methyl-2-butene (≥99.0%, Thermo Scientific), and 2-methyl-1-butene (≥98%, Thermo Scientific) were purchased and stored at –4°C until use. High-purity gases, including helium (>99.9999%), nitrogen (>99.99%), an N /CO mixture (75% N, 25% CO), and zero-grade air, were obtained from Airgas (Radnor, PA, USA). Standard mixtures for the meta-transcriptomic experiments were prepared in 120 mL serum flasks containing 70 mL of anaerobic minimal medium. The standards for the VFA exclusion experiments were prepared in 25 mL Balco tubes with 15 mL of mineral medium. Isoprene was introduced from stock solutions using gas-tight glass syringes. Standards of 3-methyl-1-butene, 2-methyl-2-butene, and 2-methyl-1-butene were prepared separately in the same manner. All gases were delivered via appropriately sized gas-tight syringes.

### Enrichment cultures

Original anaerobic enrichment cultures were established using 1.3 g of dry eucalyptus soil sediment inoculated into 120 mL sealed serum flasks containing 70 mL of sulfate-free, anaerobic minimal medium, as described by Widdel and Bak (1992). The anaerobic minimal medium contained MgCl2 x 6H2O (0.4 g/L), KBr (0.09 g/L), KCl (0.66 g/L), and CaCl2 x 2H2O (0.1 g/L). The medium was dispensed into 120 mL culture flasks, crimp-sealed with Teflon-faced rubber septa, and autoclaved. The autoclaved medium was degassed with N2 for 20 min in the medium and 15 min in the headspace, followed by degassing with 80/20 CO2/N2 for 15 min in the medium and 10 min in the headspace. Following degassing, NaHCO3 (1M, 30 mL/L), NH4Cl (5 g/L)/KH2PO4 (4 g/L) mixture (30 mL/L), trace element mixture (1 mL; nitriloacetic acid, 1.5 g/L; MgSO4 x 7H2O, 3 g/L; NaCl, 1 g/L; CaCl2 x 2H2O, 0.1 g/L; KAI(SO4)2 x 12 H2O, 0.02 g/L; Na2SeO3 x 5H2O, 0.3 mg/L; Na2WO4 x 2H2O, 0.4 mg/L; FeSO4 x 7H2O, 0.1 g/L; MnSO4 x H2O, 0.5 g/L; CoSO4 x 6H2O, 0.18 g/L; CuSO4 x 5H2O, 0.01 g/L; NiCl2 x 6H2O, 0.03 g/L; Na2MoO4, 0.01 g/L; ZnSO4 x 7H2O, 0.18 g/L; H3BO3, 0.01 g/L; distilled water, 1 L), and 1 mL vitamin solution (biotin, 2 mg/L; thiamine hydrochloride, 5 mg/L; pyridoxine hydrochloride, 10 mg/L; folic acid, 2 mg/L; riboflavin, 5 mg/L; lipoic acid, 5 mg/L; D-Ca-pantothenate, 5 mg; vitamin B12, 0.1 mg/L; 4-aminobenzoic acid, 5 mg/L; nicotinic acid, 5 mg/L; distilled water, 1 L). Vitamins and trace element solutions were filter-sterilized, and all other solutions were autoclaved.

The basal medium was supplemented with a racemic mixture of D/L-lactate and a mixture of volatile fatty acids (VFAs; acetate, butyrate, propionate, succinate, and formate) to a final concentration of 10 mM each. Isoprene (≥99.9% purity) was added as the terminal electron acceptor by injecting 0.05 mL of liquid isoprene into the bottles, yielding a headspace concentration of approximately 100–300 μmol.

Cultures were incubated under strictly anoxic conditions and subjected to sequential dilution-to-extinction enrichment after an initial 1-year incubation. Transfers were performed at a 1:1 dilution ratio and repeated eight times. The resulting eighth-generation enrichment culture was then used as the inoculum for meta-transcriptomic analyses and volatile fatty acid exclusion experiments.

### Gas Chromatopgraphy analysis

Headspace gas samples (100 μL) were collected weekly from the enrichment cultures using a pressure-lockable gas-tight syringe (Fisher Scientific, UK) and analyzed by gas chromatography (TRACE™ GC-1600). Separation of isoprene and methylbutenes was achieved on a GasPro PLOT column (60 m × 0.32 mm, Agilent Technologies) with nitrogen as the carrier gas at 5 mL/min. Detection was performed with a flame ionization detector (FID). The GC oven was programmed to hold at 50°C for 30 seconds, then increase at 20°C/min to 250°C. Manual injection of the collected headspace samples ensured consistent analysis of substrate and product concentrations.

### DNA extraction, 16S rRNA sequencing, and analysis

Genomic DNA was extracted from 1 mL of culture at time zero and after six weeks of incubation for each treatment group using the FastDNA™ Spin Kit for Soil (MP Biomedicals, USA) following the manufacturer’s protocol. Extracted DNA was stored at −80 °C until analysis. The V4 region of the 16S rRNA gene was amplified with modified Earth Microbiome primers 515F and 806R, which target both bacteria and archaea (Caporaso et al., 2011; Parada et al., 2016). Forward primers included unique barcodes, and PCR products were purified with AMPure XP beads (Beckman Coulter, USA) according to Illumina’s MiSeq workflow. Sequencing was performed on an Illumina MiSeq platform (Illumina, USA) with paired-end chemistry.

Sequence data were processed with Mothur (Schloss et al., 2009) on the Pete Supercomputing Facility at Oklahoma State University. Paired-end reads were merged into contigs, and low-quality, anomalous, and duplicate sequences were removed. Sequences were aligned against the SILVA nr_v138_2 reference database (SILVA, 2023), and poorly aligned reads and overhangs were trimmed. Chimeric sequences were identified and excluded, and unique sequences differing by ≤2 bp were clustered to reduce sequencing error. Taxonomic classification was assigned at the genus level, and non-bacterial/archaeal lineages (eukaryotic or unidentified) were removed.

### Experimental design of the VFA exclusion study

For VFA test I, the enriched inoculum from the 8^th^ dilution of a culture enriched with isoprene and the 5 VFAs (acetate, formate, butyrate, succinate, propionate) and D/L lactate (ICE1_D8) served as the original inoculum source. The experimental design consisted of 9 treatments, each conducted in biological duplicate, resulting in 18 experimental units.

Each culture was prepared in 25 mL sterile Balch tubes, with a total working volume of 15 mL, comprising 10 mL of pre-reduced anoxic mineral medium, 5 mL of enrichment inoculum (ICE1_D8), and a remaining headspace filled with an N:CO (80:20) gas mixture. Tubes were sealed with butyl rubber stoppers and aluminum crimps to maintain strict anoxic conditions.

A defined set of substrates, including lactate, fatty acids (VFAs) such as acetate, succinate, propionate, butyrate, and formate, was used as carbon sources, each added at a final concentration of 10 mM to initially enrich ICE1_D8 (inoculum). To determine the role of individual carbon sources in sustaining isoprene reduction, each treatment excluded one VFA. An additional treatment (ICE-ALL) received all carbon sources, and two controls were included: (i) a no-inoculum control (IC_Control) with all carbon sources to test for abiotic isoprene reduction, and (ii) an enrichment inoculum control (EI_Control) with inoculum but no added carbon sources to assess carryover substrates. For VFA Test II, three experimental groups were established: Treatment I: Inoculum + Formate (20 mM) + Isoprene; Treatment II: Inoculum + Formate (10 mM) + Lactate (10 mM) + Propionate (10 mM) + Isoprene; Treatment III: Inoculum + Lactate (10 mM) + Propionate (10 mM) + Isoprene (Table 1). Each group consisted of three biological replicates and uninoculated controls. All other experimental conditions were identical to those in VFA Test I.

### Experimental design for the metatranscriptomic study

The enriched culture from the eighth dilution of a highly active enrichment, named Eucalyptus soil sediment + Carbon sources + Isoprene_1 (ICE1), served as the inoculum for the experimental treatments of the meta transcriptomic study.

From this dilution, 25 mL of culture was transferred to each of four experimental treatments: ICE1.A1, ICE1.A2 (isoprene-amended), and EC1.A1, EC1.A2 (unamended controls). All cultures were supplied with the full carbon substrate mix (acetate, lactate, formate, butyrate, succinate, and butyrate; 10 mM each), and only ICE1 replicates received 0.05 mL of liquid isoprene, added in the same manner as during enrichment. Control treatments (EC1.A1, EC1.A2) were not amended with isoprene but were otherwise identical. Cultures were incubated at 25°C for 21 days until >50% of isoprene was reduced to 2-methyl-1-butene, at which point samples were harvested for RNA extraction and meta-transcriptomic analysis. Total RNA was extracted from each replicate using the RNeasy PowerFecal Pro Kit (Qiagen), optimized for complex environmental matrices, following the manufacturer’s instructions. The extracted RNA was sent to BMK gene for meta-transcriptomic sequencing and analysis.

### Library preparation and high-throughput sequencing

RNA integrity and quantity were assessed using a Bioanalyzer 2100 (Agilent Technologies) and a Qubit fluorometer (Thermo Fisher Scientific). Ribosomal RNA depletion and cDNA library construction were performed using standardized Illumina protocols. Libraries were sequenced on an Illumina NovaSeq 6000 platform with paired-end 150-bp reads (PE150), yielding an average of ∼22 million raw reads per sample, for a total of over 88 million reads across all conditions. Sequencing was performed by BMKGENE (Beijing, China).

### Quality filtering and assembly

Raw reads were filtered with Trimmomatic (parameters: LEADING:3 TRAILING:3 SLIDINGWINDOW: 5:20 MINLEN:50) to remove low-quality bases and adapter contamination. The resulting clean reads were quality-checked with FastQC. Reads mapping to rRNA were removed, and mRNA-enriched reads were assembled into transcript contigs with IDBA-Tran (https://github.com/loneknightpy/idba).

### ORF prediction and gene catalog construction

Assembled contigs were analyzed with Prodigal to predict open reading frames (ORFs), and the resulting sequences were filtered to retain only those ≥90 bp. Redundant ORFs across samples were removed with MMseqs2 to construct a non-redundant unigene catalog. Sequencing reads were aligned to the unigene set with Bowtie2 (v1.0), and gene-level abundance was quantified with BBMap and Samtools (v1.3.1) to compute raw read counts, FPKM values, and relative abundances.

### Functional and taxonomic annotation

Unigenes were annotated against multiple databases: 1) KEGG (Kyoto Encyclopedia of Genes and Genomes) for metabolic pathways, enzyme functions, and module participation; 2) eggNOG for orthologous groups and functional classification (COG categories: L2); 3) CAZy for carbohydrate-active enzyme families; 4) CARD and BacMet for antibiotic and biocide resistance genes; 5) VFDB and PHI-base for virulence and pathogen–host interaction factors; and 6) TCDB for membrane transport protein classification.

Taxonomic profiling was performed using the MetaPhlAn marker gene database, and functional annotation of unigenes was derived from the NCBI NR protein database. Taxonomic annotation was performed by aligning unigenes to the NCBI non-redundant (NR) protein database, with further refinement using MetaPhlAn3 for genus- and species-level classification. Krona charts were generated to visualize community composition across multiple taxonomic ranks.

### Multivariate and differential expression analyses

Gene expression differences between isoprene-supplemented and control cultures were assessed using comparative abundance analysis of annotated functional genes. Community-wide shifts were visualized using principal component analysis (PCA), principal coordinates analysis (PCoA), and non-metric multidimensional scaling (NMDS). PCA was used to explore variance in gene expression profiles across treatments, while Bray-Curtis-based PCoA was applied to assess differences in community composition at the species level. Analyses were based on relative abundance profiles of KEGG orthologs, eggNOG functions, and genus-level taxonomy. Heatmaps, Venn diagrams, and ternary plots were used to visualize shared and condition-specific expression patterns. Where applicable, the statistical significance of clustering was assessed using Adonis and ANOSIM tests.

## Results

### Volatile Fatty Acid exclusion study

Time-course measurements of isoprene (solid lines) and 2-methyl-1-butene (dotted lines) are shown for enrichment cultures lacking individual VFAs or all VFAs, compared with uninoculated controls. Panels show the exclusion of (a) acetate, (b) lactate, (c) formate, (d) propionate, (e) butyrate, (f) succinate, (g) all VFAs, and (h) no VFA addition. Data are shown as means of biological duplicates, with error bars indicating standard deviation.

To identify the volatile fatty acids (VFAs) required for isoprene reduction, exclusion assays were conducted by removing individual VFAs from the enrichment cultures and monitoring isoprene consumption and methylbutene production relative to the full VFA-amended control (Fig. 1a–h). Across all treatments, initial isoprene concentrations at time 0 ranged from ∼45 to 118 µmol/microcosm, with an overall mean of 77.57 ± 9.52 µmol/microcosm. By the end of 4 weeks, average 2-methyl-1-butene concentrations across all treatment groups reached 145.46 ± 18.44 µmol/microcosm, indicating substantial isoprene reduction.

**Figure 1.**
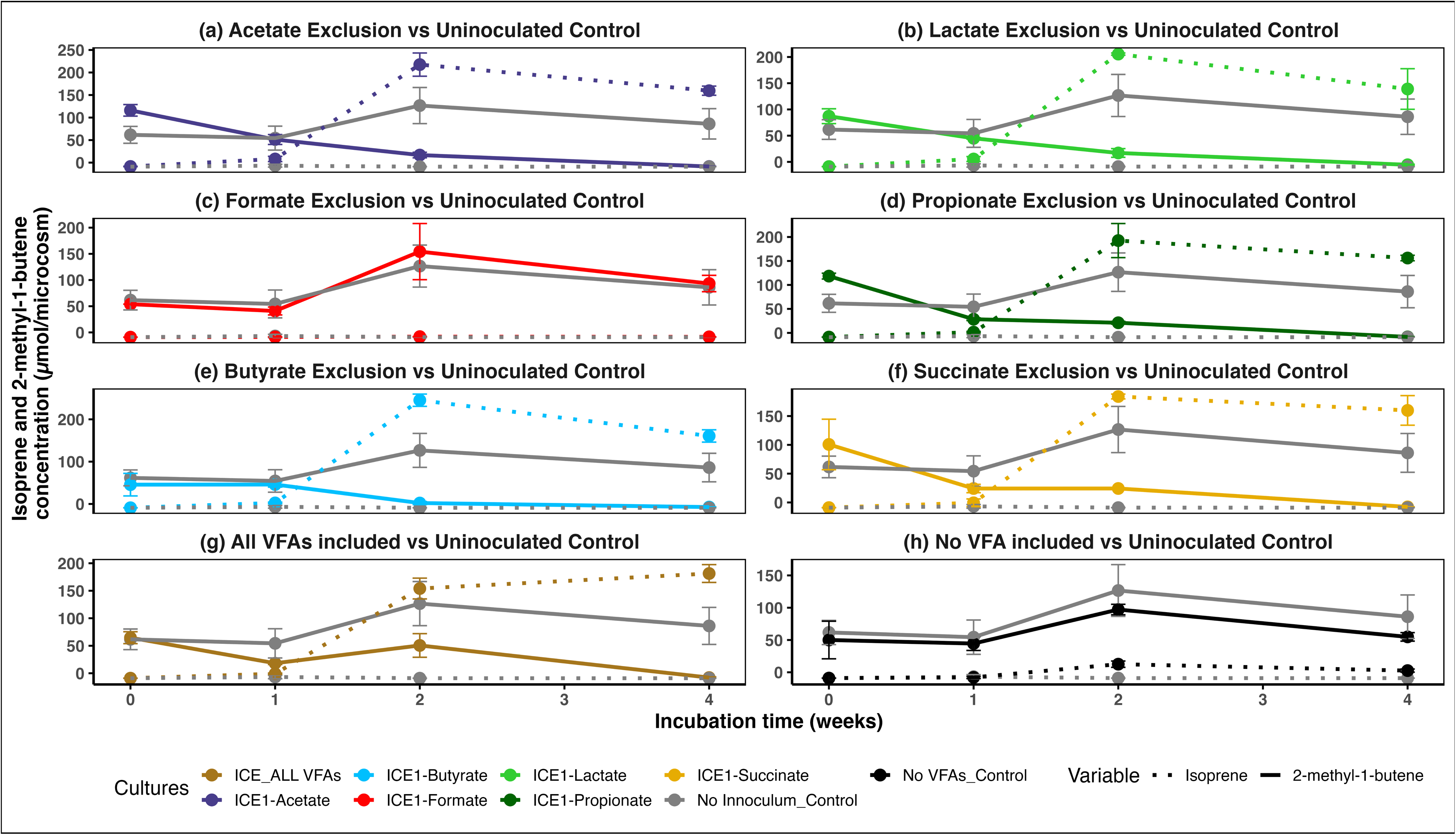
Effect of volatile fatty acid (VFA) exclusion on isoprene reduction in enrichment cultures. Time-course measurements of isoprene (solid lines) and 2-methyl-1-butene (dotted lines) are shown for enrichment cultures lacking individual VFAs or all VFAs, compared with uninoculated controls. Panels show exclusion of (a) acetate, (b) lactate, (c) formate, (d) propionate, (e) butyrate, (f) succinate, (g) all VFAs, and (h) no VFA addition. Data are shown as means of biological duplicates, with error bars indicating standard deviation.

Formate exclusion uniquely eliminated isoprene reduction. In cultures where formate was excluded (Fig. 1c), isoprene concentrations remained high throughout incubation (93.35 ± 15.60 µmol at week 4) with no 2-methyl-1-butene production. This mirrored results from the uninoculated control and the control with no added VFAs or lactate. The no-VFA inoculated control produced only 2.61 ± 2.44 µmol product and 54.84 ± 6.60 µmol isoprene (Fig. 1h), and the uninoculated control showed no product formation, with 86.06 ± 33.68 µmol isoprene remaining at week 4.

In contrast, all other VFA exclusion treatments (Fig. 1a, b, d-f) showed complete isoprene consumption and high 2-methyl-1-butene accumulation (e.g., 159.6 ± 10.1 µmol for acetate exclusion, 160.7 ± 14.6 µmol for butyrate exclusion) at the end of 4 weeks of incubation. The treatment with all VFAs and lactate introduced exhibited strong isoprene reduction activity, with isoprene fully consumed and 181.3 ± 16.2 µmol of 2-methyl-1-butene produced by week 4. Butyrate and succinate exclusions produced the highest product yields. At week 4, 2-methyl-1-butene concentrations reached 160.7 ± 14.6 µmol and 159.8 ± 25.7 µmol in butyrate- and succinate-excluded cultures, respectively (Fig. 1a, b, d–f).

Time-course data further showed that active treatments typically peaked at week 2 (mean across active panels and all-VFA, 199.8 ± 12.7 µmol) and decreased slightly by week 4 (159.4 ± 5.5 µmol), consistent with slow volatilization or redistribution of the product.

Collectively, these data demonstrate that formate is the sole non-substitutable electron donor for isoprene reduction. Excluding formate resulted in negligible 2-methyl-1-butene and sustained isoprene, whereas omitting any other VFA produced near-complete isoprene depletion and high product accumulation comparable to the all-VFA condition. Minimal production in the no-VFA and uninoculated controls further supports isoprene reduction as a formate-dependent, biologically mediated processVFA test II results

In the cultures, isoprene and formate consumption and formation of 2-methyl-1-butene were monitored in enrichment microcosms for six weeks under different substrate amendments (Fig. 2 a–f). In cultures amended with formate only (EIF, Fig. 2a), isoprene concentrations declined sharply from ∼800 µmol per microcosm at week 0 to <50 µmol by week 6 across all three replicates. The rate of isoprene loss was greatest between weeks 0 and 4, with two replicates showing >75% depletion by week 4. This rapid decrease coincided with the accumulation of the reduced product of isoprene, i.e., 2-methyl-1-butene, which reached 350–450 µmol per microcosm by the end of incubation. Uninoculated controls retained stable isoprene concentrations (600–650 µmol) throughout the incubation, with no detectable formation of 2-methyl-1-butene. Formate concentrations declined steadily in all EIF cultures over the incubation period (Figure 2b). Initial formate concentrations of approximately 14–17 mM decreased to ≤1 mM by week 6 in two replicates (EIF_2, EIF_3), while a third replicate retained low residual formate (EIF_1). In contrast, formate concentrations in uninoculated controls remained relatively constant, indicating that formate consumption was biologically mediated. In cultures amended with formate, propionate, and lactate (FLIP, Fig. 2a), initial isoprene concentrations were ∼900 µmol per microcosm and decreased variably across replicates. By week 6, one replicate retained ∼400 µmol, while another dropped below 200 µmol, indicating heterogeneous activity. The reduced product of isoprene, 2-methyl-1-butene, was not detected until week 4 and remained low, with final concentrations ranging from 50–120 µmol. Compared to the formate-only treatment, both the extent of isoprene depletion and the magnitude of product accumulation were substantially lower. Formate concentrations in lactate-, propionate-, and formate-amended FLIP cultures ranged from approximately 13–17 mM at the start of incubation (week 0) and exhibited variable trajectories across replicates over time (Figure 2d). In two replicates (FLIP_2, FLIP_3), formate concentrations fell to <5 mM by the end of 6 weeks, whereas one replicate (FLIP_1) showed no formate consumption, with a trend mirroring that of the uninoculated control. FLIP_1 is the only replicate in this treatment group that did not show detectable formation of the reduced product of isoprene, 2-methyl-1-butene.

**Figure 2.**
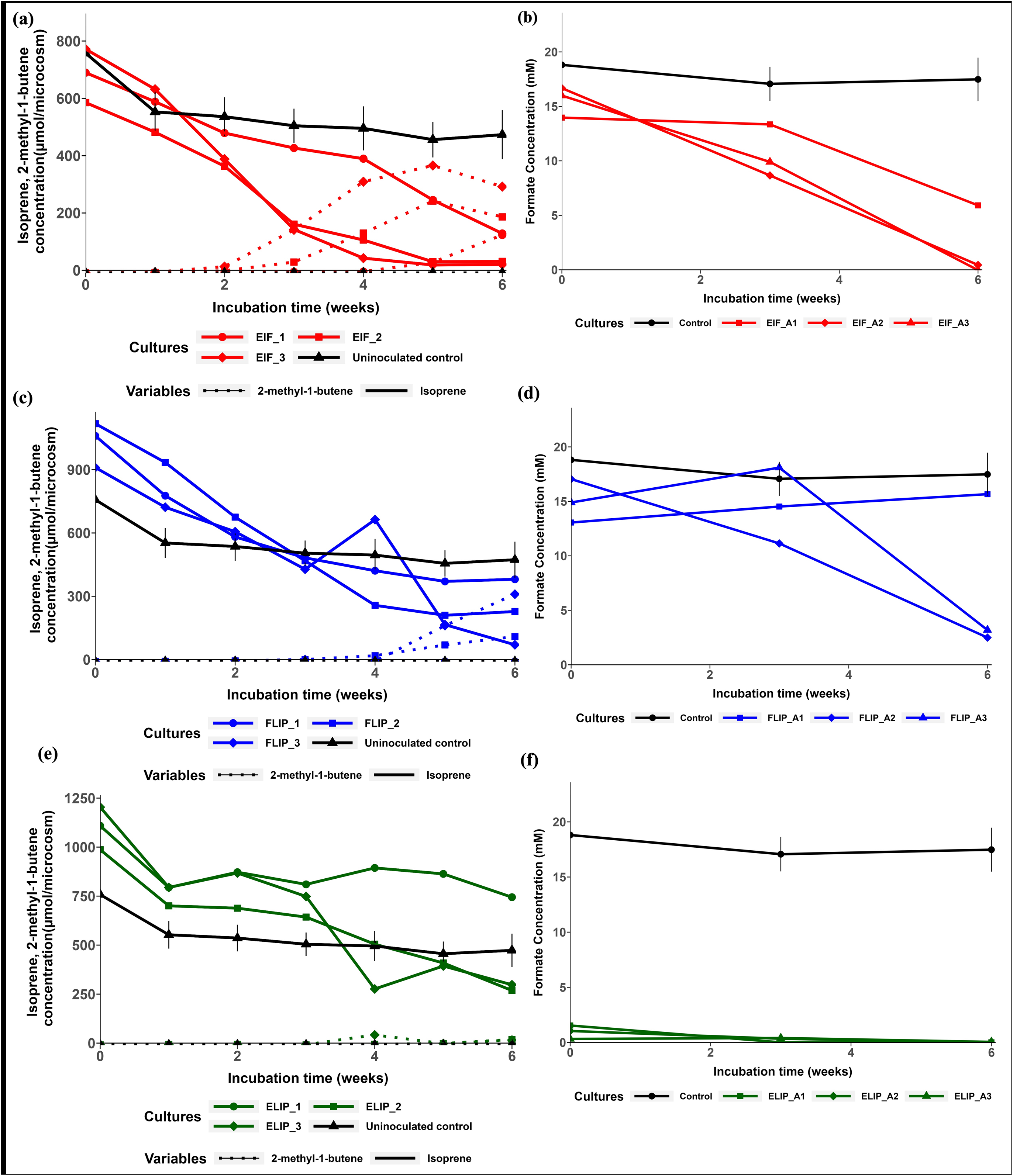
Consumption of isoprene and formate and production of 2-methyl-1-butene in enrichment cultures. Microcosms were incubated with isoprene under various substrate conditions for 6 weeks. (a, b) Inoculum + isoprene + formate (three replicates; EIF_1, EIF_2, EIF_3), (c, d) Inoculum + isoprene + formate + lactate + propionate (three replicates; FLIP_1, FLIP_2, FLIP_3), (e, f) Inoculum + isoprene + lactate + propionate (three replicates; ELIP_1, ELIP_2, ELIP_3), and formate + isoprene (uninoculated control). Isoprene concentration is shown by solid lines, and the formation of the reduced product 2-methyl-1-butene is shown by a dotted line. Error bars indicate standard error for the uninoculated control (n=3).

In cultures amended with propionate and lactate (ELIP, Fig. 2c), isoprene concentrations were higher (∼1,100 µmol) than in other treatments. Two replicates showed a modest decline to ∼700–800 µmol by week 6, while the third exhibited a sharper decrease to ∼450 µmol, suggesting variable responsiveness to ELIP. However, 2-methyl-1-butene formation was negligible, with only trace amounts (<20 µmol) detected late in the incubation. Uninoculated controls remained steady at about ∼500 µmol of isoprene with no product formation. Formate concentrations in all replicates ranged from 0.5 mM to 1.5 mM.

Collectively, these data demonstrate that formate consistently enabled complete isoprene reduction and robust production of 2-methyl-1-butene alone. Formate accelerated isoprene loss and supported >7-fold higher accumulation of 2-methyl-1-butene than other treatments, underscoring its role as the critical electron donor driving this metabolic process. These findings indicate that formate may serve a dual role for isoprene-reducing microorganisms, acting as both an electron donor for isoprene reduction and a supplementary carbon source.

### 16S rRNA community analyses results

Time 0 analysis of the inoculum (T0_1, T0_2) showed that the starting community was dominated by *Sporomusa* (60.5 ± 6.6%, n=2). Secondary taxa each comprised ≤6–7%: *Sedimentibacter* (5.8 ± 1.6%), *Pelotomaculum* (5.8 ± 0.6%), *Lachnoclostridium* (5.0 ± 1.1%), and *Lachnospiraceae unclassified* (4.6 ± 0.8%). After six weeks of incubation, the substrate amendments strongly altered community dynamics.

In formate-only treatment (EIF), the community shifted most strongly relative to T0: *Pelotomaculum* increased to 30.5 ± 4.2% (n = 3; ∼4.6 times higher than baseline), followed by *Lachnoclostridium* (20.3 ± 2.6%) and *Sedimentibacter* (12.0 ± 2.2%), while *Sporomusa* declined to 12.1 ± 3.7%. This strong enrichment of *Pelotomaculum* directly correlates with the GC results (Fig. 2a), where formate-amended cultures showed complete isoprene depletion and the highest accumulation of 2-methyl-1-butene. The co-enrichment of *Lachnoclostridium* suggests a potential role as a secondary partner, possibly in metabolite exchange, while *Sedimentibacter* may be involved in fermenting intermediate metabolites or in metabolic cross-feeding. The roles of these microorganisms in the enrichment are not entirely clear.

In the lactate and propionate (ELIP) treatment, the community remained structurally similar to T0, with *Sporomusa* remaining dominant (47.3 ± 10.4%, *n* = 3), and with increases in *Desulfitobacterium* (20.8 ± 7.1%), *Sedimentibacter* (13.2 ± 5.2%), and the fermenter *Proteiniborus* (6.9 ± 1.9%). *Pelotomaculum* and *Lachnoclostridium* remained low (3.8 ± 1.4% and 0.19 ± 0.03%, respectively). In the absence of formate, lactate and propionate favored fermentative taxa yet failed to support *Pelotomaculum* enrichment or detectable isoprene turnover (Fig. 3, 2c).

**Figure 3.**
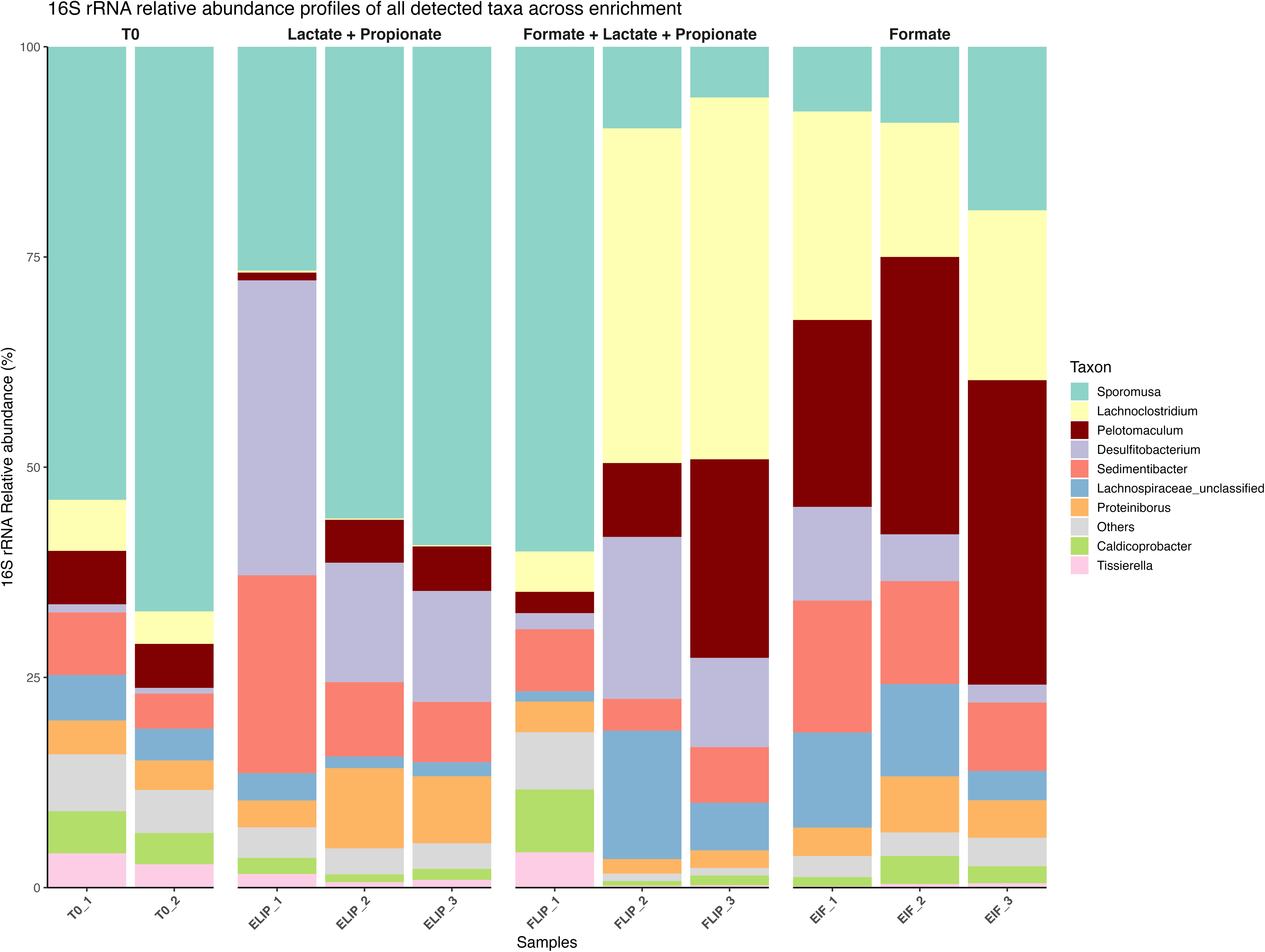
Microbial community composition across substrate amendments. Relative abundances of all detectable microbial taxa, based on 16S rRNA gene amplicon sequencing, are shown for the initial inoculum (T0) and enrichment cultures amended with lactate + propionate (ELIP), formate + lactate + propionate (FLIP), and formate alone (EIF). Bars represent biological replicates for each condition.

In the formate, lactate, and propionate (FLIP) treatment, two replicates (FLIP_2, FLIP_3) converged on a *Lachnoclostridium*-enriched state (41.42 ± 1.62%, *n* = 2), with *Pelotomaculum* at 16.20 ± 7.42% and *Sporomusa* at 7.87 ± 1.84%. In contrast, FLIP_1 diverged, remaining *Sporomusa*-dominated (60.02%) with low levels of *Pelotomaculum* (2.53%) and *Lachnoclostridium* (4.80%). This low abundance of *Pelotomaculum* likely explains why isoprene was not reduced in that replicate (Fig 2c). Although *Pelotomaculum* increased in relative abundance compared to T0, *Lachnoclostridium* was the most enriched taxon across this treatment, suggesting that competitive interactions with fermentative partners may limit the extent of isoprene reduction under mixed-substrate conditions.

### Meta-transcriptomic analysis showed shifts in the microbial community when isoprene is added

Bray–Curtis-based principal coordinate analysis (PCoA) revealed a clear, consistent separation between isoprene-amended (ICE1) and unamended control (EC1) communities (Fig. 4a). The first principal coordinate axis (PCoA1) explained 99.5% of the variation in species-level community composition, with ICE1 samples clustering tightly on the negative axis and EC1 samples on the positive axis. This pattern indicates that isoprene amendment was the dominant factor shaping microbial community structure, driving reproducible enrichment of distinct taxa in ICE1 relative to EC1. The second coordinate axis (PCoA2), which accounted for only 0.5% of the variation, captured minor within-group variability, particularly among EC1 replicates. The tight clustering of isoprene-amended replicates, contrasted with the broader dispersion of controls, indicates that isoprene exerts strong deterministic selection for a narrow consortium, whereas control communities assemble more variably through stochastic processes among alternative taxa. A normalized relative abundance heatmap was generated to compare species-level community composition between isoprene-amended (ICE1) and unamended control (EC1) cultures (Fig. 4b). The analysis shows that *Pelotomaculum schinkii* is strongly enriched in isoprene-amended replicates (ICE1.A). In contrast, replicates without isoprene (EC1.A) show higher enrichment of *Sediminibacter saalensis.* This indicates that isoprene amendment selectively suppressed fermenters while favoring *Pelotomaculum schinkii*.

**Figure 4.**
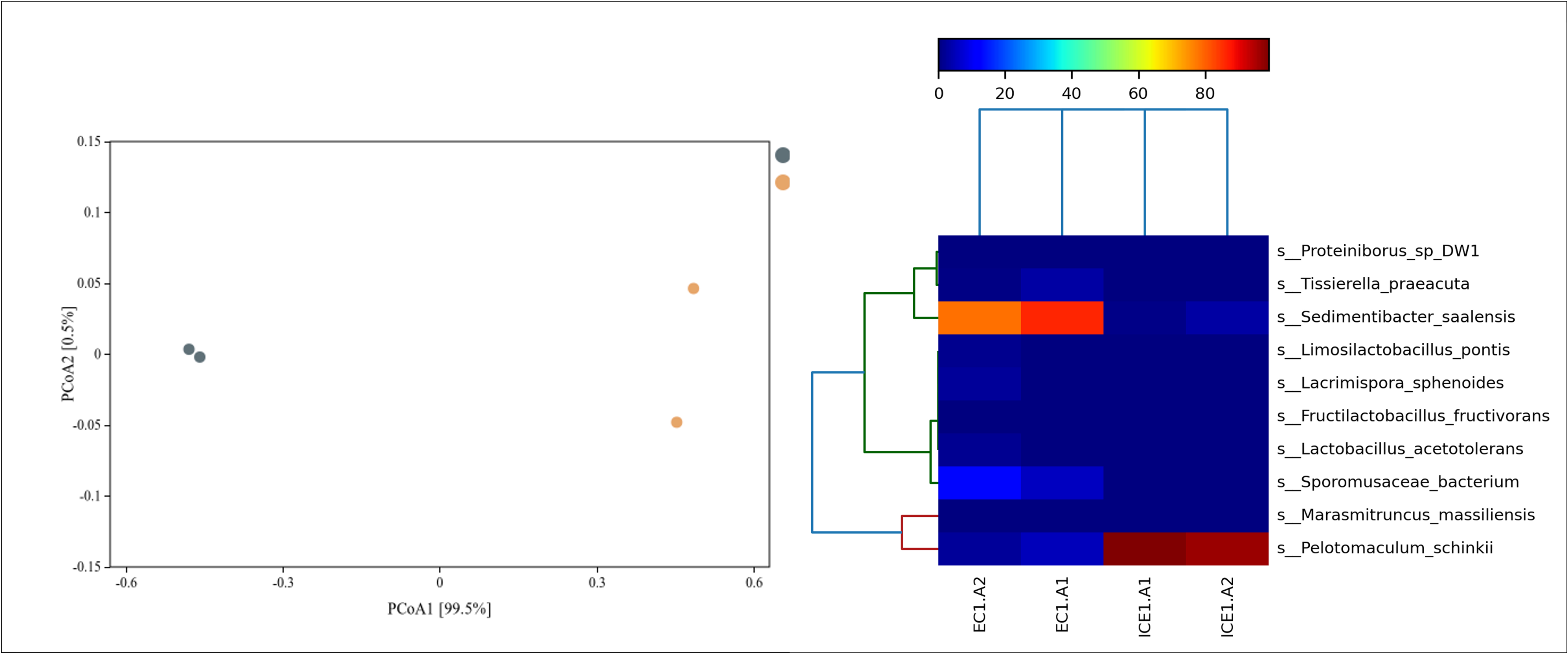
Microbial community shifts in response to isoprene amendment. **(a)** Bray-Curtis-based Principal Coordinate Analysis (PCoA) of species-level community composition in isoprene-amended (ICE1) and unamended control (EC1) cultures. **(b)** Heatmap of the top differentially expressed species across ICE1 and EC1 samples, with hierarchical clustering.

**Figure 5.**
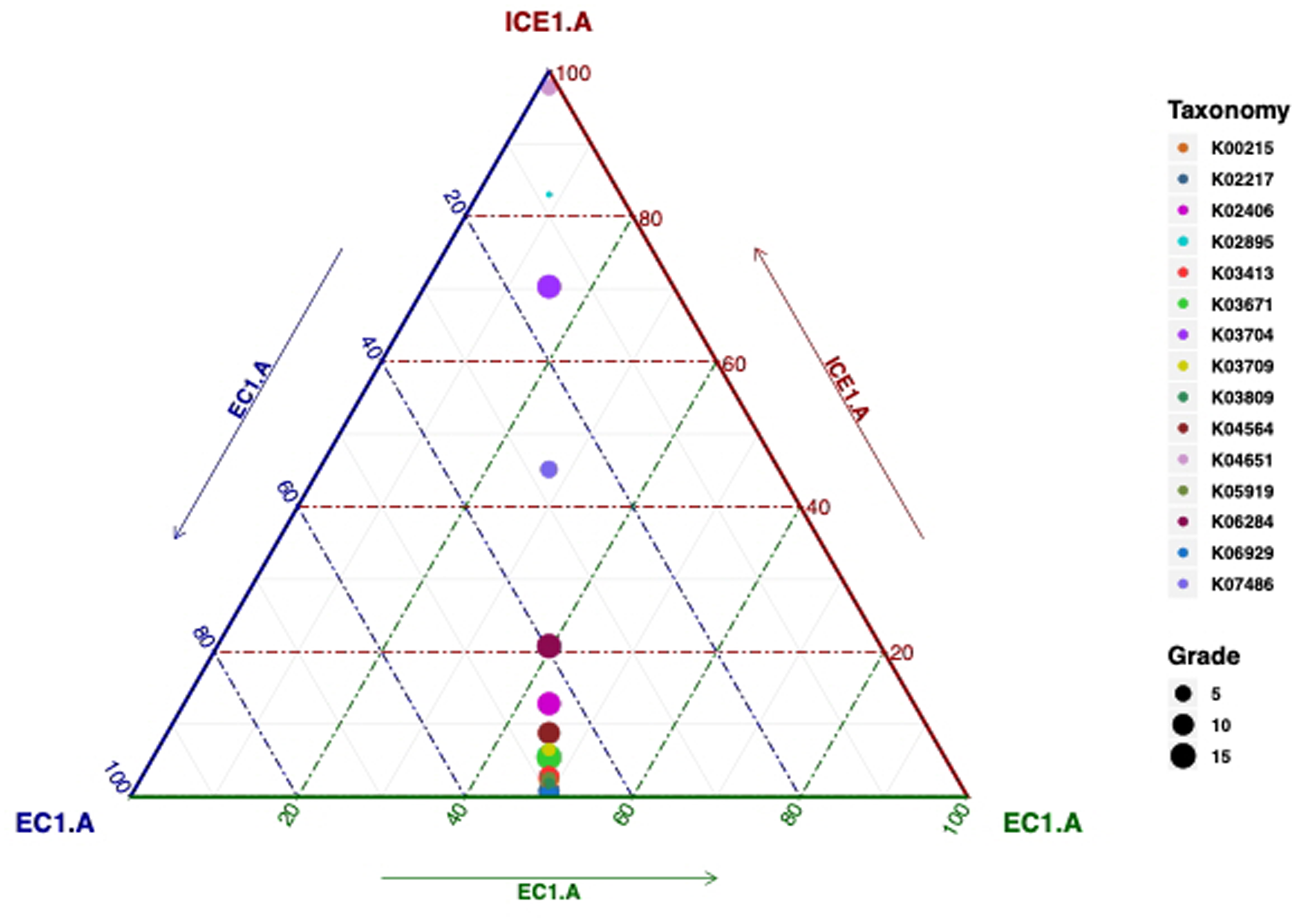
Ternary plot of differentially expressed KEGG orthologs across isoprene-reducing (ICE1.A) and control (EC1.A) conditions. Each point represents a gene, with size indicating overall expression level and color denoting KEGG ID. ICE1.A-enriched genes shift toward the top vertex, while EC1.A-enriched genes shift toward the base. Only the top 15 differentially expressed KO terms are shown.

Together, these results demonstrate that isoprene addition significantly alters the microbial community composition, selectively enriching for *P. schinkii*.

### KEGG-based ternary classification reveals functional gene enrichment in isoprene-reducing communities

Each point represents a gene, with size indicating overall expression level and color denoting KEGG ID. ICE1.A-enriched genes shift toward the top vertex, while EC1.A-enriched genes shift toward the base. Only the top 15 differentially expressed KO terms are shown. In the isoprene-amended cultures (ICE1.A), the most notable shift was the upregulation of hypA/hybF (K04651; hydrogenase nickel incorporation protein), which was nearly absent in EC1.A (0.00007) but abundant in ICE1.A (0.0064). hypA/hybF is a critical maturation factor required for nickel delivery into NiFe hydrogenases, and its selective enrichment under isoprene amendment suggests enhanced investment in hydrogenase cofactor assembly. This targeted response suggests a potential role for NiFe hydrogenase maturation in supporting electron transfer during isoprene reduction.

Other ICE1.A-enriched functions included cspA (K03704; cold shock protein, 0.0111 vs. 0.0023), indicative of a stress response to isoprene exposure, and rplX (K02895; ribosomal protein L24, 0.0049 vs. 0.0005), consistent with increased translational activity. A moderate increase in transposase K07486 (0.0033 vs. 0.0020) was also observed, suggesting genomic plasticity or mobilization of genetic elements under isoprene stress.

In contrast, isoprene-free cultures (EC1.A) showed higher transcript abundance of genes associated with redox balance, storage, and motility. These included trxA (K03671; thioredoxin, 0.0114 vs. 0.0013), consistent with enhanced redox homeostasis, and SOD2 (K04564, superoxide dismutase, 0.0047 vs. 0.0009), along with dfx (K05919; superoxide reductase, 0.0032 vs. 0.00016), highlighting a greater reliance on oxidative stress defense under control conditions. EC1.A cultures also exhibited higher expression of ferritin (K02217, 0.0051 vs. 0.00009), reflecting iron storage activity, and genes linked to chemotaxis and motility, such as cheY (K03413, 0.0039 vs. 0.0002) and flagellin fliC (K02406, 0.0053 vs. 0.0016).

Overall, isoprene amendment triggered a targeted response focused on hydrogenase maturation and stress adaptation, whereas control cultures (EC1) favored oxidative defense, nutrient storage, and motility. The selective and disproportionate enrichment of hypA/hybF in ICE1 highlights its likely importance in the physiology of isoprene reduction.

### Upregulation of metal and amino acid transporters under isoprene-reducing conditions

Each point represents a transporter classified by its **TCDB ID**, with position indicating relative expression across ICE1.A and EC1.A replicates. Transporters toward the “ICE1.A” apex are more highly expressed under isoprene-reducing conditions, while those closer to “EC1.A” are enriched in control cultures. Dot size reflects overall expression level (“Grade”) across all samples. Only the top 15 differentially expressed KO terms are shown.

Isoprene-amended cultures (ICE1.A) showed marked enrichment of several transporter families associated with ion translocation and energy conservation. A strong upregulation of H /Na - translocating F-type/V-type/A-type ATPases (3.A.2.1.14, 0.0034 vs. 0.00021; and 3.A.2.1.5, 0.0023 vs. 0.00086), along with the H /Na - translocating pyrophosphatase (M - PPase, 3.A.10.2.4, 0.0037 vs. 0.000014), was observed. These proton/sodium-translocating complexes are central to establishing ion gradients and proton motive force, suggesting enhanced energy conservation capacity under isoprene amendment. ICE1.A cultures also showed elevated transcription of the NADH dehydrogenase complex (NDH, 3.D.1.5.1, 0.0025 vs. 0.00099), supporting reinforced electron transport activity.

Isoprene-amended cultures also showed enriched transport functions, including the ferrous iron uptake system FeoB (9.A.8.1.14, 0.0031 vs. 0.00086), consistent with increased iron acquisition under isoprene, and the pore-forming amphipathic peptide HP2-20 family (1.C.82.1.1, 0.0029 vs. 0.00066), which may indicate stress-associated membrane remodeling. Increased expression of the nuclear mRNA exporter family (3.A.18.1.1, 0.0022 vs. 0.00094) suggests enhanced transcriptional and translational activity, supporting accelerated protein synthesis for redox-active and energy-conserving processes under isoprene amendment.

In contrast, the isoprene-free cultures (EC1.A) showed higher expression of transporters involved in nutrient uptake and redox protection. Members of the ATP-binding cassette (ABC) superfamily were dominant in EC1.A, with multiple transporters (3.A.1.15.8, 0.0089 vs. 0.00025; 3.A.1.5.19, 0.0023 vs. 0.0001) enriched relative to isoprene-amended cultures (ICE1.A). EC1.A also showed elevated abundance of the Na-transporting carboxylic acid decarboxylase (NaT-DC, 3.B.1.1.1, 0.00165 vs. 0.00014), which catalyzes the decarboxylation of organic acids (such as oxaloacetate or methylmalonyl-CoA) while simultaneously exporting sodium ions (Na) across the membrane. The higher abundance of NaT-DC in isoprene-free cultures suggests that these communities rely more heavily on fermentative or carboxylate-driven metabolism for energy generation when isoprene isn’t available as an electron sink. The nitrogen fixation complex FixABCX (3.D.12.1.1, 0.0013 vs. 0.00078), which mediates electron bifurcation, transferring electrons to nitrogenase for N fixation or related redox processes, is also upregulated in isoprene-free cultures, likely to re-oxidize reduced cofactors (e.g., NADH, ferredoxin) in the absence of isoprene as an external electron acceptor. The peroxiredoxin heme-binding protein Tpx1 (8.A.147.1.2, 0.0017 vs. 0.00005), which detoxifies peroxides and regulates heme-iron and siderophore-iron transport systems, was also more abundant in EC1.A, consistent with its role in oxidative stress defense.

Together, these findings highlight distinct transporter investment strategies between the two conditions. In the isoprene-amended cultures (ICE1.A), there was pronounced upregulation of energy-conserving ion-translocating systems, including H /Na - coupled ATPases, pyrophosphatases, and ferrous iron uptake pathways, consistent with enhanced chemiosmotic energy conservation and metal cofactor acquisition required for redox-active enzymes such as NiFe hydrogenases. In contrast, the isoprene-free cultures (EC1.A) exhibited higher expression of ABC-type transporters, Na-dependent carboxylic acid decarboxylases, and antioxidant proteins, including the hemin-binding peroxiredoxin Tpx1, reflecting a metabolic shift toward substrate-level energy generation, broad nutrient scavenging, and oxidative stress mitigation.

### Differential expression of metal resistance and stress response genes in ICE1.A and EC1.A

In the isoprene-amended cultures (ICE1.A), transcriptional responses were characterized by increased expression of genes linked to metal import and metabolic functions. Notably, the molybdenum transport system components modB (0.00068 vs. 4.9 × 10) and modC (0.00028 vs. 3.5 × 10) were strongly enriched in isoprene-amended cultures (ICE1.A). This shift suggests greater investment in molybdenum acquisition under isoprene. Elevated molybdenum uptake aligns with the requirement for several molybdenum-dependent redox enzymes, such as formate dehydrogenase, which catalyzes formate oxidation required for isoprene reduction. Additional ICE1.A-enriched genes included fabK (0.00035 vs. 0.00012), involved in fatty acid biosynthesis, and galE (0.00034 vs. 0.00013), which encodes UDP-glucose 4-epimerase, pointing to broader metabolic restructuring in the presence of isoprene.

In contrast, the isoprene-free cultures (EC1.A) showed substantially higher expression of genes associated with oxidative stress defense and heavy-metal efflux. These included arsT (0.00936 vs. 0.00035), a thioredoxin reductase within an arsenic resistance cluster, and arsB (0.00022 vs. 1.7 × 10), an arsenite efflux pump. Strong enrichment was also observed for sodA (0.0046 vs. 0.00060), a manganese-dependent superoxide dismutase, and mntA/ytgA (0.0026 vs. 4.7 × 10), which delivers manganese for antioxidant functions. Other EC1.A-enriched genes included the glycerol facilitator glpF (0.0021 vs. 1.6 × 10) and multiple metal-responsive transcriptional regulators, including csoR, corR, and smtB/ziaR.

These patterns indicate that isoprene amendment redirected transcription toward molybdenum acquisition and metabolic processes tightly linked to formate utilization and isoprene reduction. In contrast, control cultures prioritized detoxification systems and antioxidant defenses, including arsenic efflux, manganese transport, and copper/zinc homeostasis.

### Differential expression of unigenes reveals functional shifts beyond annotated pathways

Heatmap analysis of the top differentially expressed unigenes showed a strong treatment-specific separation between isoprene-amended (ICE1.A) and control (EC1.A) cultures, with replicates clustering closely within each condition. Across biological replicates, 18 unigenes were consistently upregulated in isoprene-amended cultures (ICE1). The most enriched transcripts included unigene_133152 (unannotated), unigene_116031 (unannotated), unigene_98337 (copper amine oxidase domain protein), unigene_139673 (cspA, cold-shock protein), unigene_65937 (hypothetical protein Psfp_02137), and unigene_103132 (hypothetical). Several features were uniquely detected in isoprene-amended cultures, including unigene_86605 (hypothetical), unigene_110732 (50S ribosomal protein L15), unigene_13870 (unannotated), and unigene_36685 (30S ribosomal protein S17), indicating condition-specific expression. A redox gene, unigene_131384 (pyridine nucleotide–disulfide oxidoreductase mapped to *P. propionicum*), was also notably enriched in ICE1.

Two hydrogenase maturation genes (hypA/hybF and hypB) from *Pelotomaculum schinkii* were among the most differentially upregulated in isoprene-amended cultures. Unigene_131371 (hypA/hybF) had an average expression of 0.00507 in ICE1.A, compared with 1.36 × 10 in EC1.A, corresponding to a ∼373-fold increase. Similarly, unigene_131357 (hypB) had an ICE1.A mean of 0.00516 versus 8.82 × 10 in EC1.A, yielding a ∼585-fold upregulation. Together, these results indicate that isoprene amendment induces a transcriptional program centered on NiFe-hydrogenase maturation and electron-transfer capacity, accompanied by increased investment in translation and stress adaptation.

## Discussion

### Beyond obligate syntropy: *P. schinkii* gains independence with isoprene

Microbial interactions range from loose associations to strict mutual dependence (Schink, 2002). In some cases, one partner supplies metabolites such as vitamins or amino acid precursors, allowing the other to conserve biosynthetic energy even if it retains the capacity to synthesize them. More obligate forms of cooperation are especially common among anaerobes. Although many bacteria and archaea in anaerobic communities are suspected of being in obligate syntrophic relationships, our knowledge of obligate syntrophs is limited to a few species belonging to the genera *Desulfofundulus*, *Pelotomaculum*, *Smithella*, *Syntrophobacter,* and *Syntrophobacterium* (Morris et al., 2013b).

From a thermodynamic standpoint, obligate syntrophy is one of the most energetically constrained microbial lifestyles, in which short-chain fatty acid (VFA) oxidizers operate near the thermodynamic limit of energy conservation (McInerney et al., 2009a; Morris et al., 2013b; Westerholm et al., 2021). The standard Gibbs free energy for oxidizing VFAs such as propionate (ΔG°^’^ ≈ +76.1), butyrate (ΔG°^’^ ≈ +48.6), and benzoate (ΔG°^’^ ≈ +70.1) is highly endergonic, and these reactions are only possible when coupled to the rapid removal of hydrogen or formate by partner methanogens or sulfate reducers (McInerney et al., 2009a). Even then, the Gibbs free energy available to syntrophic bacteria often hovers near the minimum required to sustain ATP synthesis (−15 to −20 kJ mol ¹) and can be as low as −10 kJ mol ¹ (Adams et al., 2006; McInerney et al., 2009b; Scholten & Conrad, 2000). These marginal conditions yield extremely slow growth rates and low biomass yields, making them difficult to culture in the lab.

Until now, *Pelotomaculum schinkii* has been described as a strictly obligate syntroph, best known for oxidizing propionate in association with methanogens (Field, de Bok et al., 2005). Propionate oxidation is highly endergonic under standard conditions (ΔG°′ = +73.8 kJ mol ¹ propionate) and can proceed only when hydrogen and formate concentrations are kept extremely low (≤2.85 Pa H or ≤1.58 μM formate) (Field, de Bok et al., 2005; Hidalgo Ahumada et al., 2018; Thauer et al., 1977). However, our results show that *P. schinkii* can sustain growth and conserve energy independently when isoprene is available as a terminal electron acceptor.

In our enrichment cultures, methane formation, methanogens, and sulfate reducers were undetectable, yet isoprene reduction occurred consistently, and *P. schinkii* remained the most enriched species under isoprene-amended conditions. Rather than collapsing without a syntrophic partner, *P. schinkii* maintained a high relative abundance, indicating that isoprene serves as an alternative electron sink, allowing it to decouple from classical syntrophic dependence. Meta-transcriptomic data further confirmed *P. schinkii* as the primary organism responsible for isoprene reduction: in isoprene-amended microcosms (ICE1), *P. schinkii* dominated, whereas unamended controls (EC1) were enriched in *Sedimentibacter saalensis (Fig 4b).* Ordination and hierarchical clustering of transcriptomes revealed clear separation between treatments, with *P. schinkii* enriched exclusively under isoprene-reducing conditions (Fig 4b). This shifts the view of *P. schinkii* from an obligate syntroph to a facultative anaerobe capable of opportunistic respiration using unconventional electron acceptors such as isoprene. Such metabolic flexibility likely confers a selective advantage in anoxic environments where formate and volatile hydrocarbons, including isoprene, are abundant.

### Functional and genetic evidence for formate-driven isoprene respiration

Substrate exclusion experiments demonstrated that formate is the critical electron donor supporting isoprene reduction (Fig. 1). Cultures lacking formate showed no isoprene reduction (Fig. 1c, 2e) and no enrichment of *Pelotomaculum* (Fig. 3), whereas formate-amended treatments exhibited rapid isoprene depletion, accumulation of methyl butene (Fig. 1(a-b, d-g), Fig. 2a, c), and a marked increase in *Pelotomaculum* relative abundance from Day 0 to 42 weeks (Fig. 3). This consistent pattern across enrichment assays establishes formate as indispensable for directing electron flow into the isoprene-reducing pathway. Formate serves as a versatile electron donor in diverse inducible respiratory chains across bacteria and archaea. Its low redox potential (E°′ for the CO /HCOO couple ≈ −0.43 V) enables coupling to a broad spectrum of terminal electron acceptors, including nitrate, sulfate, fumarate, Fe³, arsenate, and even dioxygen (Maia et al., 2015). For example, *Dehalococcoides mccartyi* used formate as an efficient electron donor for reductive dichlorination of trichloroethene (TCE), achieving faster and more efficient dichlorination than lactate or citrate (Tomita et al., 2022). Similarly, enrichment cultures of the anaerobic methanotrophic archaeon *Candidatus Methanoperedens nitroreducens* were recently shown to couple formate oxidation to nitrate reduction, with upregulation of formate dehydrogenase genes and proteins (Xie et al., 2023). In classical syntrophic systems, however, formate exhibits concentration-dependent effects: at low concentrations (<10 mM), formate stimulates syntrophic propionate oxidation by enhancing acetate production and promoting the growth of syntrophic partners, whereas higher concentrations inhibit metabolism by downregulating formate dehydrogenase and hydrogenase transcription and blocking the methylmalonyl-CoA pathway (Li et al., 2024). Together, these studies reinforce the view that formate serves as a versatile electron donor across phylogenetically and functionally distinct anaerobes, a role echoed in our findings, where *the microcosm* relied exclusively on formate to sustain isoprene reduction.

In isoprene-amended cultures (ICE1.A, Fig. 6), we observed exclusive activation of several energy-conserving systems, including pyrophosphate-energized proton pumps (H-PPase), which couple PPi hydrolysis to proton translocation across membranes (Baltscheffsky et al., 1999), and Na-translocating pyrophosphatases, which use the same mechanism to drive sodium ion translocation (Luoto et al., 2011; Malinen et al., 2007). Additional upregulated functions included the H-translocating ATP synthase and ferrous iron uptake (feoB).

**Figure 6.**
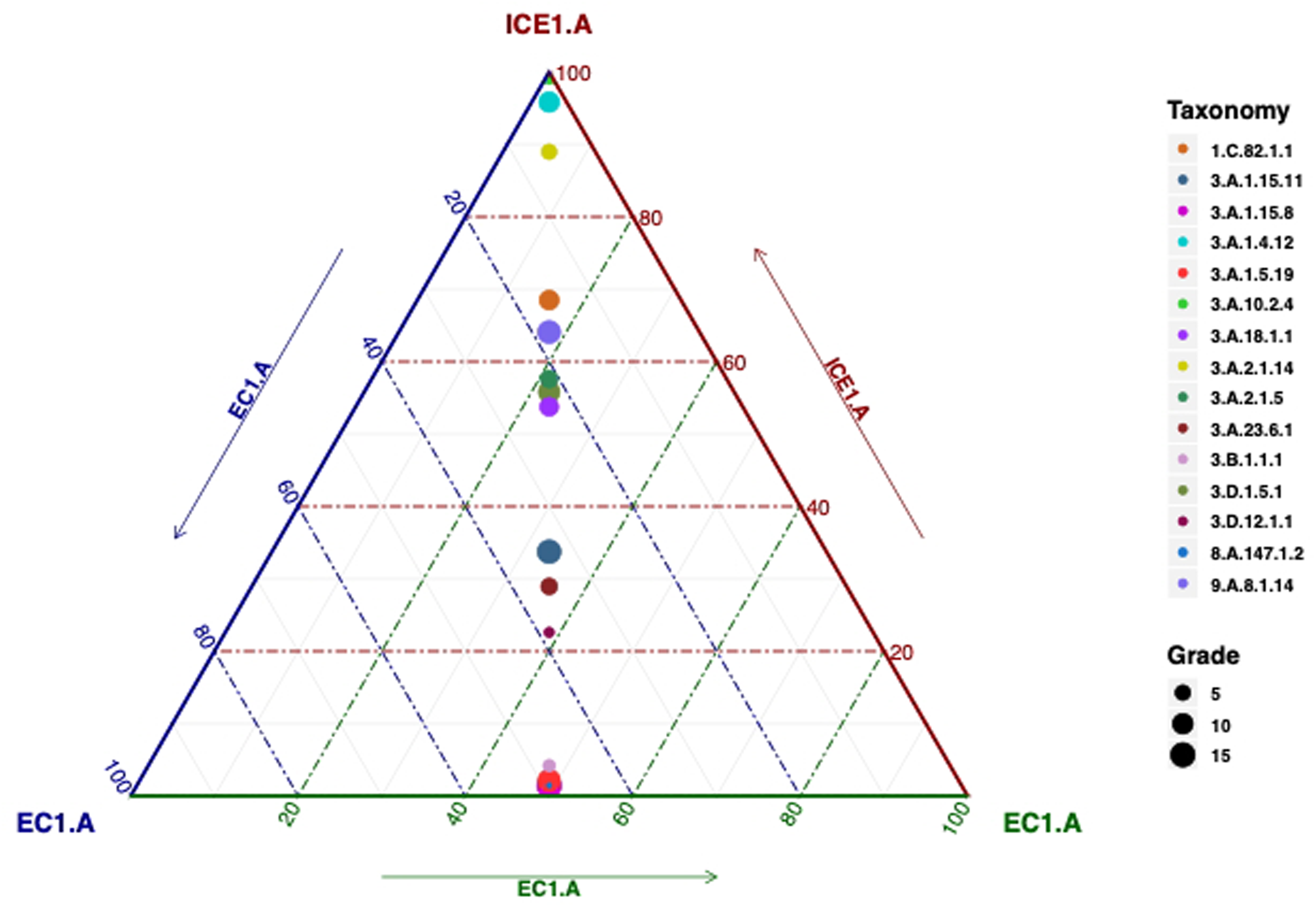
Ternary plot showing differential expression of transporter families (TCDB classification) between isoprene-reducing (ICE1.A) and control (EC1.A) conditions. Each point represents a transporter classified by its **TCDB ID**, with its position indicating relative expression in ICE1.A and EC1.A replicates. Transporters near the “ICE1.A” apex are more highly expressed under isoprene-reducing conditions, whereas those closer to “EC1.A” are enriched in control cultures. Dot size reflects overall expression level (“Grade”) across all samples. Only the top 15 differentially expressed KO terms are shown.

Genes of the molybdate transport system were also strongly induced, with modB and modC upregulated by about 14- and 8-fold, respectively, in isoprene-amended cultures (ICE1) compared to isoprene-free controls (Fig. 7). Molybdenum is a critical metal cofactor required for the catalytic activity of redox enzymes such as formate dehydrogenases (Adamus et al., 2024; Maia et al., 2015). Although components of formate dehydrogenase were not among the differentially expressed genes highlighted in the heatmap or ternary plots, direct mapping of transcripts against the *P. schinkii* strain JJ genome revealed transcriptional upregulation of genes associated with *formate dehydrogenase (NAD) activity* (GO:0008863), *formate dehydrogenase complex* (GO:0009326), and *formate metabolic process* (GO:0015942), consistent with our physiological evidence that formate serves as the primary electron donor for isoprene reduction. In contrast, transcripts associated with folate-linked C assimilation, including methylenetetrahydrofolate dehydrogenase (NADP) activity (GO: 0004488) and methylenetetrahydrofolate reductase (NADPH) activity (GO:0004489), were comparatively downregulated in isoprene-amended cultures. Consistent with the downregulation of folate-linked C enzymes, several GO categories associated with C - dependent biosynthesis were also significantly downregulated in the isoprene-amended condition, including purine nucleotide biosynthetic and metabolic processes (GO:0006164, GO:0006163), methionine metabolism and biosynthesis (GO:0006555, GO:0009086), S-adenosylmethionine (SAM)-dependent methyltransferase activity and SAM binding (GO:0008757, GO:1904047), as well as glycyl-tRNA aminoacylation and serine O-acetyltransferase activity (GO:0004820, GO:0006426, GO:0009001). Together, these patterns indicate reduced flux of formate-derived C units into nucleotide, amino acid, and methylation pathways in *P. schinkii* under isoprene-amended conditions. This can be interpreted as a shift away from C-driven anabolism toward predominantly dissimilatory formate oxidation, in which formate is exploited mainly as an electron donor to support isoprene reduction and energy conservation, while carbon for biomass is supplied by co-amended VFAs (e.g., acetate and propionate). In isoprene-free controls, higher expression of C folate enzymes likely reflects greater reliance on formate-derived one-carbon units for biosynthesis and redox balancing in the absence of an additional electron sink. This supports the involvement of the formate oxidation machinery in the active redox network during isoprene reduction. A complete list of upregulated and downregulated genes of *P. schinkii* identified through GO enrichment analysis is provided in the Supplementary Information (Table S5, S6).

**Figure 7.**
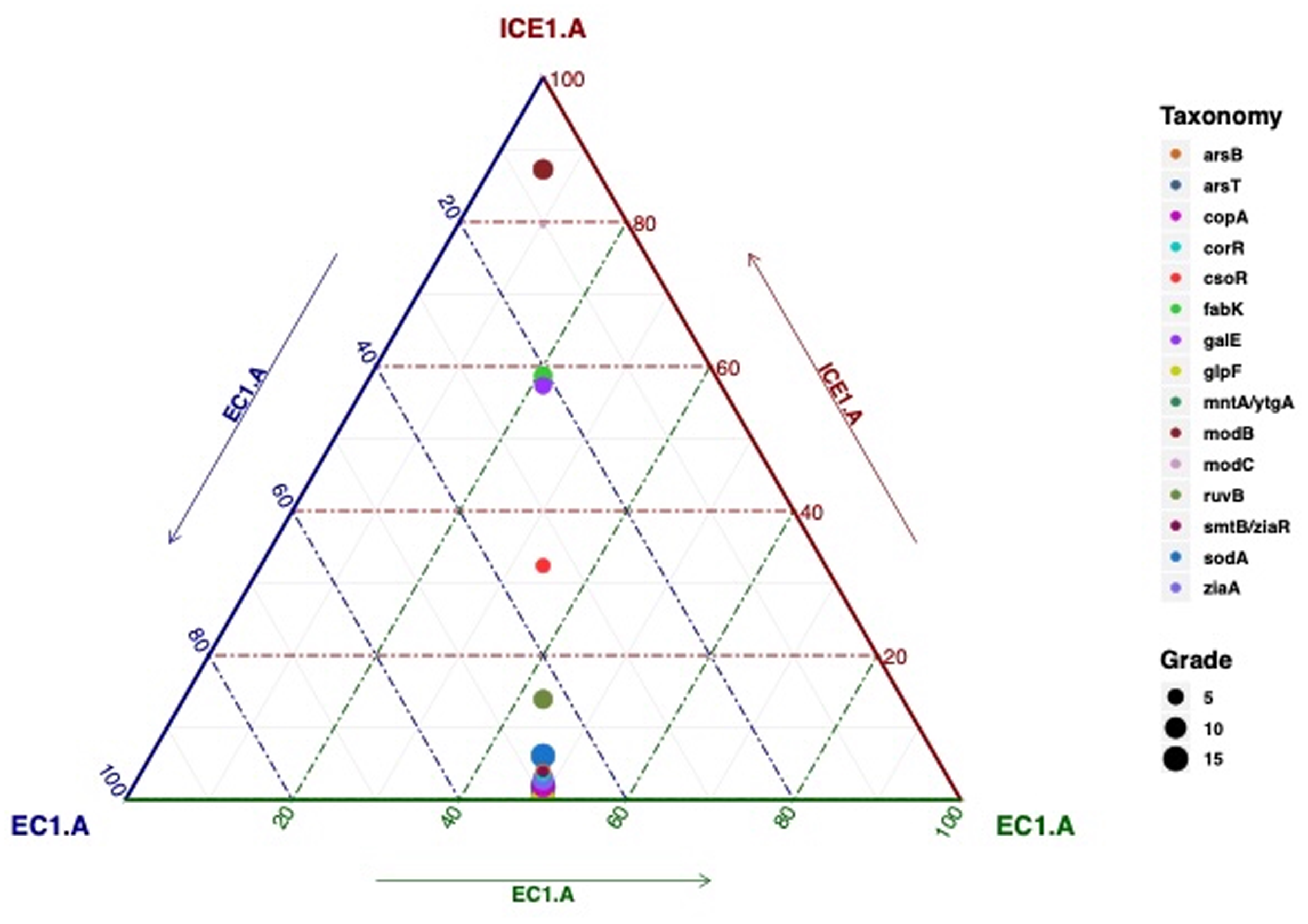
Ternary plot of metal resistance and detoxification gene expression between isoprene-amended (ICE1.A) and control (EC1.A) cultures. Each point represents an individual gene (colored by gene name, right legend) and is positioned according to relative transcript abundance across treatments. Circle size corresponds to expression grade (5–15, bottom legend). Only the top 15 differentially expressed KO terms are shown.

The heatmap (Fig. 8) showed that transcripts for the hydrogenase maturation factors HypA/HybF (unigene_131371) and HypB (unigene_131357) were enriched by about 373- and 585-fold, respectively, in isoprene-amended cultures relative to controls, and both mapped to *P. schinkii*. These transcriptional signatures parallel observations in *Acetobacterium wieringae*, where Kronen et al. (2023) identified hypA and hypB within the isr operon despite the absence of structural [NiFe]-hydrogenases, suggesting a specialized role for these maturation factors in assembling a Ni-dependent isoprene reductase (IsrA).

**Figure 8.**
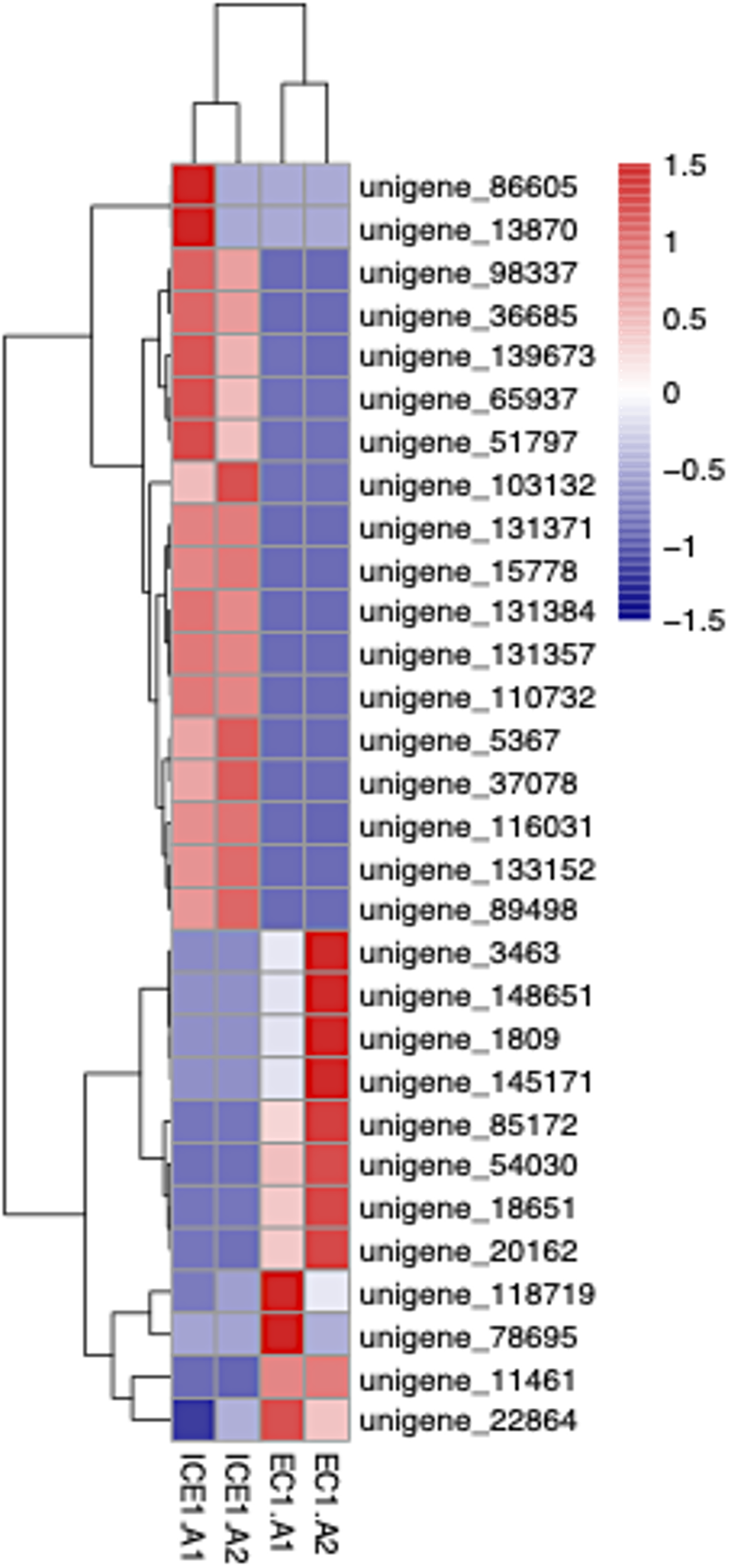
Heatmap of differentially expressed unigenes between isoprene-amended (ICE1.A) and control (EC1.A) conditions. Expression values are log-transformed and z-score normalized across samples. Genes highlighted in red are upregulated, and those in blue are downregulated. The top 30 differentially regulated unigenes are shown here.

In contrast to *Acetabacterium wieringae*, *Pelotomaculum* spp.—including *P. thermopropionicum*, *P. propionicum*, and *P. schinkii*—encode canonical [NiFe]-hydrogenases of group 1a. However, only *P. schinkii* has a group 4 (4b and 4e) NiFe hydrogenase (Westerholm et al., 2021). Group 4 [NiFe]-hydrogenases are considered among the most ancient hydrogenases, share structural similarity with the Respiratory Complex I/NADH: ubiquinone oxidoreductase in eukaryotes, and are thought to have originated in alkaline hydrothermal systems where nickel, iron, sulfur, and natural proton gradients could drive the earliest chemiosmotic reactions (Finney & Sargent, 2019, p. 2016; Russell & Nitschke, 2017; Søndergaard et al., 2016). Their coupling to simple electron donors such as formate and ferredoxin suggests they were central to primordial energy metabolism (Baradaran et al., 2013). Group 4 [NiFe]-hydrogenases can be further classified into 9 subgroups (A-I). Members of group 4b are involved in formate- and/or CO-dependent hydrogen production and are predicted to have 13 subunits, of which 7 are MRP-like antiporter subunits that generate sodium motive force linked to proton reduction (Finney & Sargent, 2019). Members of this group 4b are also well known for their ability to grow on formate as the sole carbon and energy source (Burger et al., 2022; Kim et al., 2010; Lim et al., 2014). One example is *Thermococcus onnurineus*, a hyperthermophilic archaeon in which a group 4b [NiFe]-hydrogenase (Mfh2–Mnh2) couples directly with a formate dehydrogenase (Fdh2) to support formate-driven growth (Kim et al., 2010). In this system, energy conservation is achieved through formate-dependent sodium ion pumping mediated by the Mnh2 MRP-type subunits of the group 4 hydrogenase, which in turn drives sodium-dependent ATP synthesis (Lim et al., 2014). The genome of *Thermococcus onnurineus* NA1 encodes multiple copies of formate dehydrogenases and hydrogenases, with three distinct *fdhA* genes (48–61% amino acid identity) distributed across separate gene clusters(Field, Lee et al., 2008). Of eight hydrogenase clusters, three occur in close proximity to *fdh* genes, forming modules that combine formate dehydrogenases, membrane-bound multimeric [NiFe]-hydrogenases, and cation/proton antiporters(Field, Kim et al., 2010). A similar gene cluster that integrates multiple formate dehydrogenases with [NiFe]-hydrogenase, accessory maturation proteins, and ion-translocating subunits is also found in the genome of *P. schinkii* strain JJ available in the NCBI database (Fig. 9), directly linking formate oxidation to membrane bioenergetics(Field, Hidalgo Ahumada et al., 2018).

The gene cluster in *P. schinkii* strain JJ includes accessory genes for formate dehydrogenase (FDH) assembly, including fdhE, followed by twin-arginine translocation (Tat) components (tatAd, tatC2). The Tat system allows the FDH enzyme complex to acquire cofactor before being transported across the cytoplasmic membrane (Palmer & Berks, 2012). *P. schinkii* encodes a complete suite of FDH catalytic (FdhA, FdnG), electron-transfer (FdnH, FdoH), and maturation (FdhE) proteins, representing both cytoplasmic and membrane-bound FDH systems. Together, these enzymes allow formate to be oxidized through multiple parallel pathways. Cytoplasmic FdhA-type FDHs transfer electrons directly into intracellular redox pools such as reduced ferredoxin, NADH, or F H, which supply reducing equivalents for central carbon assimilation pathways, including the generation of C metabolites like 5,10-methylene-THF, glycine, and serine (Maia et al., 2015). At the same time, the presence of FdnGHI- and FdoGHI-like complexes provides membrane-associated routes for formate-derived electrons to enter the electron transport chain via multi-heme cytochromes and Fe-S proteins, thereby generating an ion motive force (Maia et al., 2015). Interleaved with these are group 4 [NiFe] hydrogenase modules, the maturation protease hybD, and HypB nickel-incorporation proteins required for assembly of the Ni–Fe active site.

The cluster also includes an MRP-type antiporter, predicted to couple redox reactions to Na /H translocation, and a periplasmic NiFeSe hydrogenase, a high-turnover enzyme capable of reversible H oxidation/evolution (Finney & Sargent, 2019). Together, these features form a tightly integrated module in which electrons from FDH are funneled via Fe–S proteins and cytochromes into the group 4 hydrogenase, driving chemiosmotic ion translocation and enabling ATP synthesis. The embedded NiFeSe hydrogenase acts as a bidirectional redox buffer, catalyzing reversible H evolution or uptake in response to intracellular redox pressure. Excess reducing equivalents generated during formate oxidation can be dissipated via proton reduction, whereas under oxidizing conditions, H oxidation regenerates reduced cofactors. This dynamic coupling provides a ‘safety valve’ that stabilizes electron flow and maintains redox homeostasis.

This genomic architecture of *P. shinkii*, with co-localized FDHs, group 4 [NiFe]-hydrogenases, maturation factors, Tat export machinery, and an MRP-type antiporter, provides a mechanistic basis for coupling formate oxidation to energy conservation. It also aligns with our experimental findings: formate was the only electron donor required for isoprene reduction; *P. schinkii* was transcriptionally enriched in isoprene-amended cultures; and meta-transcriptomic data showed strong upregulation of [NiFe]-hydrogenase maturation factors (hypA/F, hypB), molybdate transporters (modB, modC), and chemiosmotic energy-conserving systems (ATP synthase, H - and Na - translocating pyrophosphatases). Together, these results support a model in which *P. schinkii* directly couples formate oxidation to isoprene reduction, conserving energy through ion-motive force generation and ATP synthesis via a membrane-bound formate-oxidizing NiFe hydrogenase.

### The Two Possible Mechanistic Models of Formate-Driven Isoprene Reduction

Physiological experiments (Figs. 4.1 & 4.2) demonstrate a formate-dependent isoprene reduction system, likely mediated by a group 4b [NiFe]-hydrogenase. In this system, the formate dehydrogenase complex oxidizes formate, releasing electrons that are transferred via Fe–S clusters to the [NiFe]-hydrogenase. Concomitantly, MRP-type antiporters are predicted to translocate sodium ions across the membrane, thereby generating an ion motive force (IMF). Transcriptomic data show strong upregulation of ATP synthase and H /Na-translocating pyrophosphatases in isoprene-amended cultures (ICE1), suggesting that this IMF is harnessed for ATP synthesis.

Thermodynamic considerations further support the feasibility of this metabolism. Formate oxidation coupled to proton reduction yields negligible free energy (ΔG°′ ≈ –1.3 kJ mol ¹), whereas coupling formate oxidation to isoprene reduction is predicted to release approximately –135.7 kJ mol ¹ (Kronen et al., 2023). This substantial energetic difference underscores the ecological advantage of linking formate oxidation to isoprene reduction in anoxic environments.

#### 1. Formate oxidation coupled with isoprene reduction

2HCOO^−^ ⟶ 2CO2 + 2H^+^ + 2e^−^, ΔG^0^ = −1.3 kJ/mol

C_5_H_8_ + 2H^+^ + 2e^−^ ⟶ C_5_H_10_, ΔG^0^ = −137 KJ/mol (Kronen et al, 2023)

#### Net Reaction

HCOO^−^ + C_5_H_8_ + H^+^ ⟶ CO_2_ + C_5_H_10_, ΔG^0^ = ∼ −135.7 KJ/mol

Based on these findings, two mechanistic models can be proposed:

Model A: Direct Reduction by [NiFe]-Hydrogenase: In this model, the [NiFe]-hydrogenase directly reduces isoprene by accepting electrons from formate oxidation and catalyzing the conversion of C H to C H.

Model B: Indirect Reduction via IsrA-like Oxidoreductase: Alternatively, the [NiFe]-hydrogenase may operate in a canonical manner, releasing hydrogen that is then utilized by an isrA-like oxidoreductase (analogous to the one found in *Acetobacterium wieringae*). In this scenario, isoprene reduction occurs secondarily via hydrogen-dependent catalysis.

Together, these models raise a critical mechanistic question: Does *P. schinkii* use [NiFe]-hydrogenase as a direct isoprene reductase, or rely on hydrogen-mediated transfer to an isrA-like oxidoreductase? Resolving this distinction will be essential to understanding the biochemical basis of formate-driven isoprene respiration.

## Conclusions

This study provides the first integrated evidence that *Pelotomaculum schinkii* can couple formate oxidation to isoprene reduction, thereby escaping its classical dependence on syntrophic partners. In enrichment cultures, *P. schinkii* thrived without methanogens or sulfate reducers, with formate as the sole indispensable electron donor. Physiological, transcriptomic, and genomic evidence converge on a formate-driven respiratory pathway in which electrons from formate dehydrogenase are transferred via group 4b [NiFe]-hydrogenases and associated electron carriers, and ion-translocating antiporters and ATP-generating complexes couple this process to chemiosmotic energy conservation.

Two mechanistic models emerge: (i) direct isoprene reduction by the group 4b [NiFe]-hydrogenase, or (ii) indirect reduction mediated by hydrogen produced from the hydrogenase and consumed by an isrA-like oxidoreductase. Both scenarios represent a novel respiratory mode in *P. schinkii*, broadening its ecological potential beyond classical syntrophic metabolism.

Future work should focus on resolving this mechanistic ambiguity by isolating pure isoprene-reducing cultures, conducting proteogenomic analysis, biochemically characterizing group 4b [NiFe]-hydrogenase and isrA-like proteins, and performing targeted inhibition studies. At the ecosystem scale, examining the distribution and regulation of this pathway across anoxic environments will clarify the role of formate-dependent isoprene respiration as a previously unrecognized sink in microbial carbon cycling.

## Acknowledgement

This study was supported by start-up funding provided to Dr. Sabrina Beckmann by the Department of Microbiology and Molecular Genetics at Oklahoma State University.

## Data Availability Statement

In this study, amplicon sequencing data used for community analysis are deposited in the NCBI Sequence Read Archive (SRA) under BioProject PRJNA1153434. 16S rRNA sequence data for samples are available under BioSample SUB15781987, and the raw meta-transcriptomics sequence data are available under BioSample SUB15784900.

---

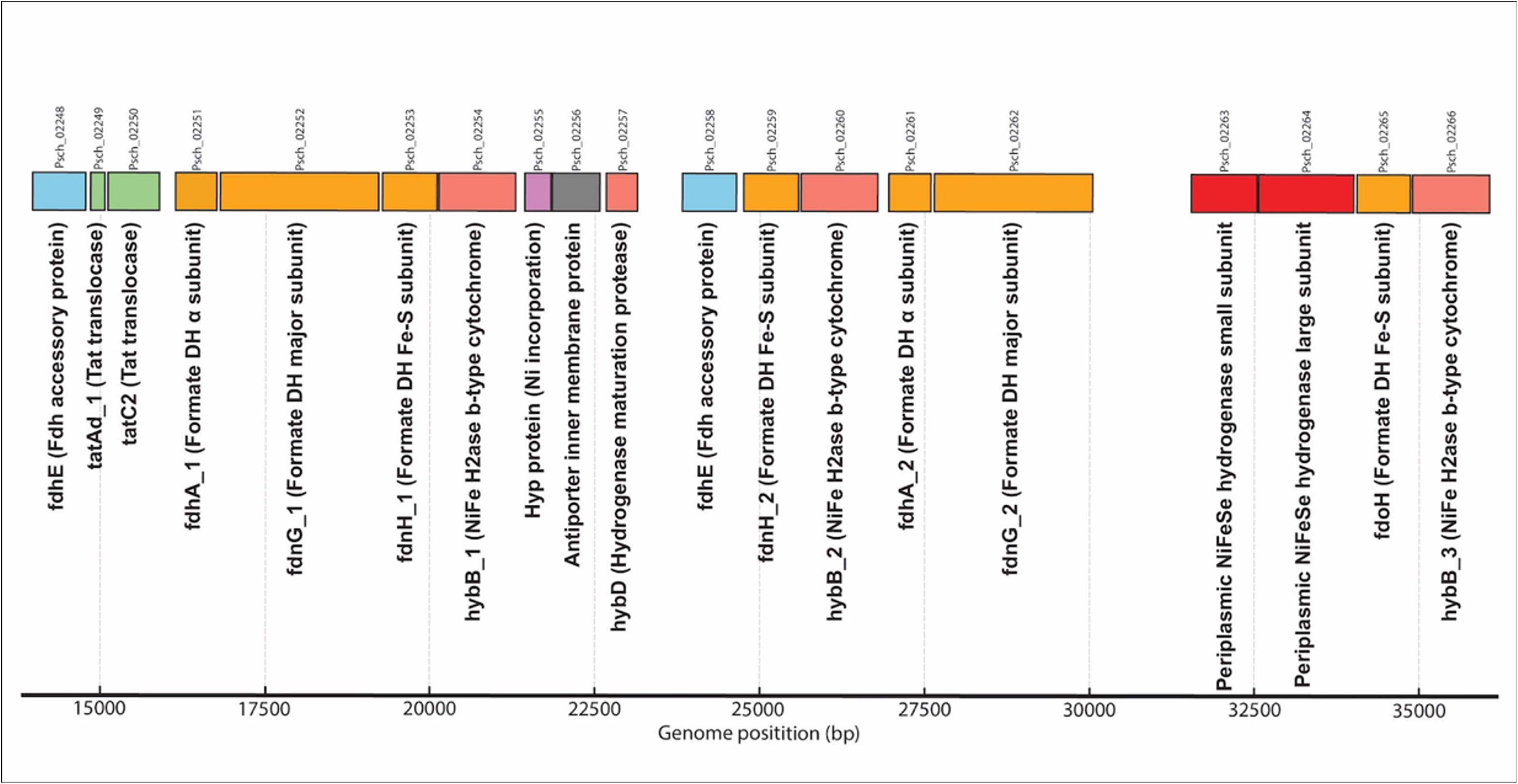

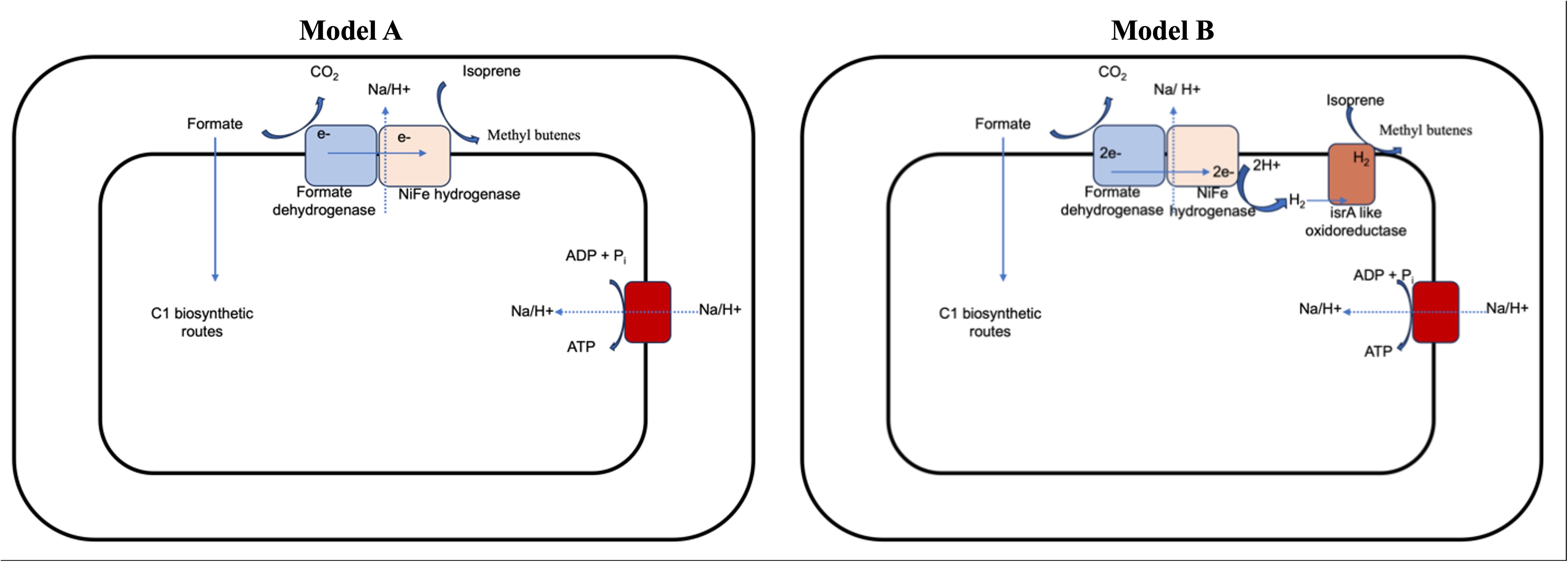

